# Ecological networks: Pursuing the shortest path, however narrow and crooked

**DOI:** 10.1101/475715

**Authors:** Andrea Costa, Ana M. Martín González, Katell Guizien, Andrea M. Doglioli, José María Gómez, Anne A. Petrenko, Stefano Allesina

## Abstract

Representing data as networks cuts across all sub-disciplines in ecology and evolutionary biology. Besides providing a compact representation of the interconnections between agents, network analysis allows the identification of especially important nodes, according to various metrics that often rely on the calculation of the shortest paths connecting any two nodes. While the interpretation of a shortest paths is straightforward in binary, unweighted networks, whenever weights are reported, the calculation could yield unexpected results. We analyzed 129 studies of ecological networks published in the last decade and making use of shortest paths, and discovered a methodological inaccuracy related to the edge weights used to calculate shortest paths (and related centrality measures), particularly in interaction networks. Specifically, 49% of the studies do not report sufficient information on the calculation to allow their replication, and 61% of the studies on weighted networks may contain errors in how shortest paths are calculated. Using toy models and empirical ecological data, we show how to transform the data prior to calculation and illustrate the pitfalls that need to be avoided. We conclude by proposing a five-point check-list to foster best-practices in the calculation and reporting of centrality measures in ecology and evolution studies.

## INTRODUCTION

The last two decades have witnessed an exponential increase in the use of graph analysis in ecological and conservation studies (see refs 1&2 for recent introductions to network theory in ecology and evolution). Networks (graphs) represent agents as nodes linked by edges representing pairwise relationships. For instance, a food web can be represented as a network of species (nodes) and their feeding relationships (edges)^3^. Similarly, the spatial dynamics of a metapopulation can be analyzed by connecting the patches of suitable habitat (nodes) with edges measuring dispersal between patches^4^. Data might either simply report the presence/absence of an edge (binary, unweighted networks), or provide a strength for each edge (weighted networks). In turn, these weights can represent a variety of ecologically-relevant quantities, depending on the system being described. For instance, edge weights can quantify interaction frequency (e.g., visitation networks^5^), interaction strength (e.g., per-capita effect of one species on the growth rate of another^3^), carbon-flow between trophic levels^6^, genetic similarity^7^, niche overlap (e.g., number of shared resources between two species^8^), affinity^9^, dispersal probabilities (e.g., the rate at which individuals of a population move between patches^10^), cost of dispersal between patches (e.g., resistance^11^), etc.

Despite such large variety of ecological network representations, a common task is the identification of nodes of high importance, such as keystone species in a food web, patches acting as stepping stones in a dispersal network, or genes with pleiotropic effects. The identification of important nodes is typically accomplished through centrality measures^5,12^. A large number of centrality measures has been proposed, each probing complementary aspects of node-to-node relationships^13^. For instance, Closeness centrality^14,15^ highlights nodes that are “near” to all other nodes in the network in terms of average distance (calculated as number of edges) from all other nodes. Whenever the effects of a node on another weaken along the path^16^, then central nodes are those having the largest capacity to influence the others. Consider however highly modular networks, in which tightly knit communities of nodes are loosely connected to one another; then, one may be interested in identifying nodes that act as bridges connecting the different communities, allowing for the spread of perturbations across the entire network. Stress centrality^17^, and Betweenness centrality^15^ serve this purpose. The choice of a centrality measure thus depends on the research question at hand, and on the characteristics of the data being analyzed. Different centrality measures have been used to identify keystone species in networks of biotic interactions^5,18^, to explore the robustness of metapopulations^19^, to describe connectivity patterns across fragmented habitats^20^, to explore social behavior and pathogen spread within populations^21^, and to provide a theoretical background to support decision-making in conservation planning and urban management^11,22^ (see Supporting Information for a complete list). Given the wide array of available techniques and the span of ecological applications, confusion may arise when performing and reporting centrality analysis. An understanding of how the calculations are performed, as well as a clear and sufficient reporting of the details of the analysis, are necessary in order to avoid misinterpretation of the results and ensure the reproducibility of published studies.

In particular, edge weights exert a substantial influence on all measures of centrality. Many centrality metrics rely on calculating the shortest path connecting any two nodes, and for weighted edges this translates into finding the path with the smallest sum of weights. Edge weights definition is crucial in all measures of centrality. When edge weights represent cost, resistance, or in general scale inversely to the strength of the relationship between two nodes, then the definition of “shortest” paths retains its simple interpretation. However, whenever edge weights are proportional to the strength of the relationship between two nodes (e.g., probability of dispersal, interaction frequency, contact rate, etc.), then minimizing the sum of edges along the path makes no sense: the data need to be transformed prior to the analysis, or one has to choose an appropriate method able to deal with this situation. This issue has been raised before^23,24^ and is widely acknowledged among studies describing community structure, where measures of network structure have been specifically developed to account for weighted edges^25–27^. However, our analysis of the literature suggests that this issue is not fully resolved, and that incorrect interpretations of centrality measures linger on, particularly among weighted networks.

Furthermore, most published studies do not report with sufficient detail the calculation of the node-to-node distance definition used in conjunction with the calculated centrality measures. As a consequence, it is often impossible to evaluate the correctness of the calculations. To quantify the extent of this problematic, we performed a systematic analysis of the ecological network literature. We selected all ecological studies from the Web of Science that use a network approach, by searching on topics TS=(network AND ecolog*), and limiting our search to Articles written in English in the science-related citation indexes (SCI-EXPANDED, CPCI-S, BKCI-S, ESCI) since 2006. We further added 22 records obtained from other sources. From the resulting list of articles, we refined our search to those mentioning “centrality”, and from this final list of 210 articles we selected studies that studied ecological communities using centrality metrics requiring the calculation of shortest paths, discarding purely methodological studies. Finally, armed with a list of 129 articles, we checked whether the analysis was reported with sufficient detail to determine whether the calculations were appropriate. In Figure 1 we summarize this information with a frequency chart and in the Supporting Information we provide the full list of articles. 63 articles (49%) did not report sufficient information on the calculation of centrality to allow their replication, six of which were unclear even whether edges were binary or weighted. Moreover, 61% of the studies using weighted edges may contain errors in how shortest path centralities are calculated, a figure that grows to 89% if we limit the analysis to the case of weighted interaction networks. Noticeably, 88% of the studies that correctly accounted for weighted edges in the calculation of shortest paths, considered networks in which weights are inversely proportional to the strength of association – thus not requiring transformation. Furthermore, nine studies (eight of which examine interaction networks) calculated centrality using the binary version of the weighted data, without providing any justification for this methodological choice. Certain ecological sub-disciplines use alternative metrics for the study of energy flow across networks, such as food-chain metrics to describe energy flow along food webs. Such metrics operate differently than shortest-path centralities, and are not reviewed in this article.

**Figure 1.**
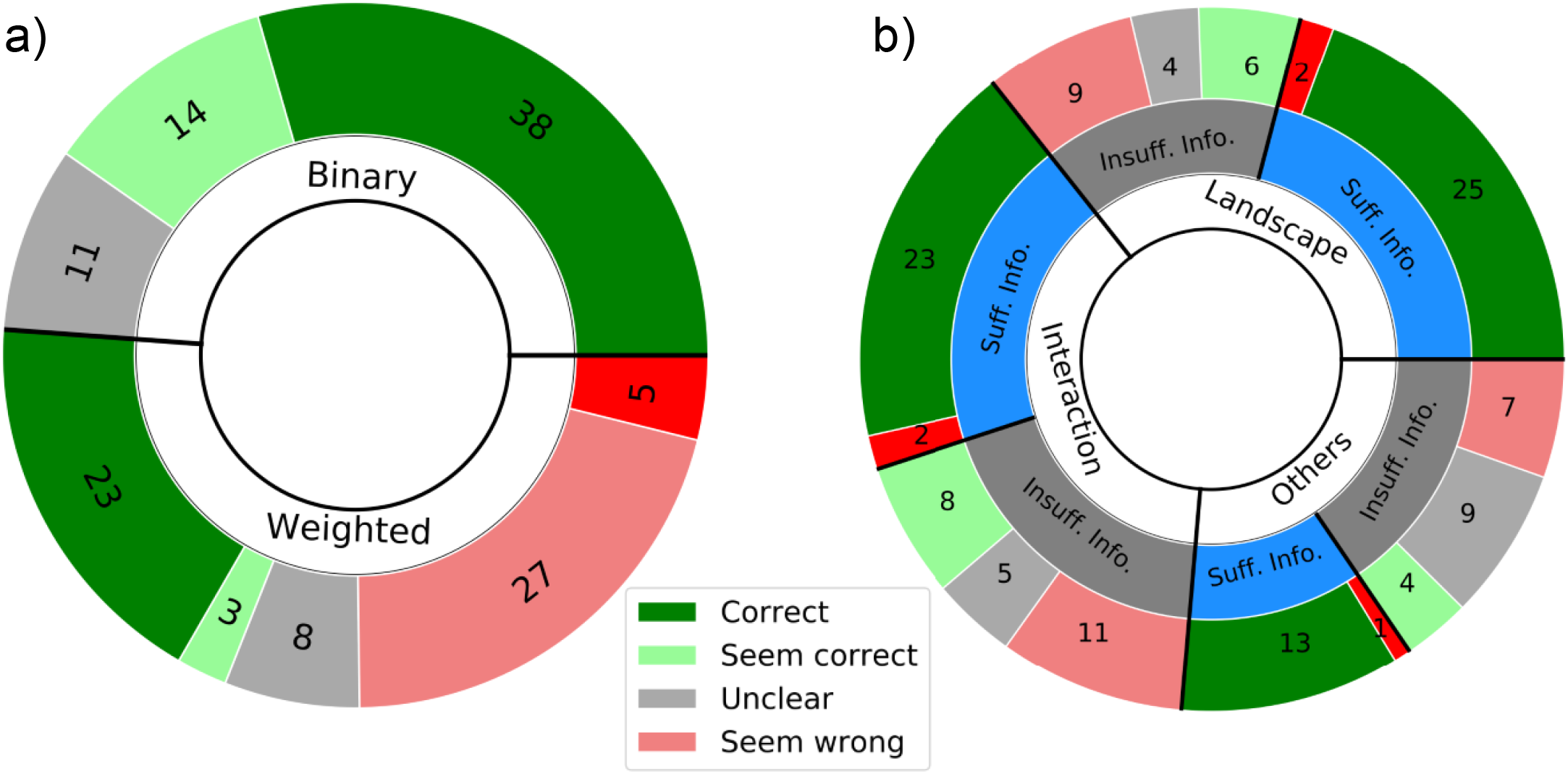
Summary of our literature analysis, detailing the number of studies by (a) edge weights, and (b) whether the information provided was sufficient or insufficient, and network type (as “landscape”, “interaction”, and “others”, which include social, co-occurrence, etc. networks). We only categorize as “correct” or as “wrong” studies on which we had enough information to support such a claim, and as “seems correct” and “seems wrong” studies on which the insufficient information available suggest that calculations are correct or wrong, respectively. Notice that (1) most of the correct or probably correct studies use unweighted edges (n = 52, 66%); (1) for all network types half of the studies do not report enough information to validate whether calculations were correct (n = 63, 49%); and that (3) numerous studies report unclear calculations (n = 19). Differences in the number of oversights in centrality calculations between weighted and landscape and interaction networks are probably due to the fact that in interaction networks weighted edges typically require transformation (see Table 1), whereas in landscape networks edge weights tend to be inversely proportional, requiring no transformation.

The final goal of this study is to offer a precise and detailed protocol for the calculation of shortest path centralities in ecological networks. Given that the correct way to calculate shortest path-based centrality measures depends on the type of network considered (binary or weighted) and on the meaning of edge weights, we use simple examples as well as real data to show how different approaches can result in unreliable estimates of centrality. Finally, since most studies omit to report key aspects of the definition of the edge weights, we propose a simple checklist to foster best-practices in the calculation and reporting of shortest path-related centrality analysis.

**Table 1.** Summary of different network types, describing the type of information flowing along edges and the consequences for weight transformation. Landscape networks are spatially-explicit, while interaction networks depict relationships among entities. Social networks are here included as interaction networks, where shared traits include shared interacting organisms, foraging time, etc. depicting social relationships or common characteristics.

| Network type | Edge weight | Proportionality to<br>information flow | Requires<br>transformation | Example<br>references |
| --- | --- | --- | --- | --- |
| Landscape | Cost-distance | Inverse | No | 11, 20 |
|  | Dispersal<br>probability | Direct | Yes | 10, 41 |
|  | Dispersal time | Inverse | No | 12 |
|  | Exchange of<br>individuals | Direct | Yes | 42 |
|  | Genetic similarity | Direct | Yes | 43, 44 |
| Interaction | Frequencies | Direct | Yes | 45, 46 |
|  | Shared traits,<br>affinity | Direct | Yes | 47, 48 |
|  | Trophic (or energy)<br>flow | Direct | Yes | 49, 50 |

## THE SHORTEST PATH IS FULL OF PITFALLS

In network analysis, the interaction between nodes can be thought of as a flow of information between the nodes that are linked by edges. The sequence of edges that information must cross in order to reach a specific node is called a path. It is generally assumed that the bulk of information between any two given nodes (among all the possible paths between these two nodes) passes through the shortest path connecting them (i.e., the one with “lowest weight”). However, it should be emphasized that while the concept of information flow is general, its immanence can differ dramatically from case to case, depending on which network feature weights quantify. For reference, in Table 1 we list the principal types of networks and the proportionality of the edges to the information flow between nodes found in the ecological literature.

The interpretation of a shortest path as the path that funnels the bulk of information flow relies on it being the least weight path (i.e., the path of least resistance) between two nodes. Indeed, all the shortest path algorithms currently available^28,29^ and generally implemented in graph theory software (Table 2) seek to minimize the value of the path between two nodes calculated as the sum of the edge weights. The reason being that a minimization problem converges, while maximization can fail (see ref 24 for a detailed explanation). Nevertheless, the identification of the shortest paths is far from trivial, as one must pay attention to what edge weights represent. That is, one has to ensure that the edge weight is inversely proportional to the flow of information between the nodes. This condition is automatically fulfilled if the natural weight suggested by the network at study is already inversely proportional to the information flow (e.g., resistance distance, dispersal time). However, when the weight is directly proportional to the information flow (e.g., interaction frequency, individual transfer, dispersal probability, pathogen transmission, energy transfer across food webs), it is necessary to transform the edge weight in order to calculate the shortest paths, and the centrality measures that rely on them (Table 3). In particular, this is important when using user-friendly software packages that automate the calculation of centrality measures (Table 2).

**Table 2.** Analytical packages commonly used for the calculation of centrality in ecological studies and references for each package. Only *sna* and *Ucinet* packages caution in their documentation about edge transformations when calculating shortest paths.

| Package | References |
| --- | --- |
| igraph | 51, http://igraph.org/ |
| sna | 52, https://cran.r-project.org/web/packages/sna/index.html |
| tnet | 53, https://cran.r-project.org/web/packages/tnet/index.html |
| Pajek | 54, http://mrvar.fdv.uni-lj.si/pajek/ |
| Ucinet | 55, https://sites.google.com/site/ucinetsoftware/home |
| CONEFOR | 56, http://www.conefor.org/ |
| Graphab | 57, https://sourcesup.renater.fr/graphab/en/home.html |

**Table 3.** Measures of shortest path-related centrality measures commonly used in ecological network analysis. Notice that other common centrality measures are based on eigenvectors or dissimilarity scores instead of on the identification of shortest paths and are hence not considered in this work.

| Centrality Measure | Definition | Formula | Intended Network type | Reference |
| --- | --- | --- | --- | --- |
| Betweenness, BC | Quantifies the proportion of shortest paths $g$ between any two nodes $i, j$, that pass through a focal node $v$. | $BC(v) = \frac{\sum_{i \neq v \neq j} g_{ij}(v)}{g_{ij}}$ | All types | 15 |
| Stress, SC | Measures the number of shortest paths $g$ between any two nodes $i, j$, that pass through a focal node $v$. | $SC(v) = \sum_{i \neq v \neq j} g_{ij}(v)$ | All types | 17 |
| Closeness, CC | Measures the average length of the shortest paths from a node $v$ to all the other nodes in the network. | $CC(v) = \frac{N-1}{\sum_j g_{vj}}$ | All types | 15 |
| Integral Index of Connectivity, IIC | Measures the degree of connectivity of the entire landscape (of total area $A_L$) through the calculation of the number of edges in the shortest path $nl_{ij}$ between patches with area $a_i$ and $a_j$. | $IIC(v) = \frac{\sum_{i=1}^n \sum_{j=1}^n a_i a_j / (1 + nl_{ij})}{A_L^2}$ | Binary landscape networks | 58 |
| Probability of Connectivity Index, PC | Quantifies the probability that two species randomly placed across a patchy landscape (of total area $A_L$) fall into habitat patches $a_i$ and $a_j$ that are reachable from each other with a maximum connectivity probability $p_{ij}$, defined as the maximum product probability of all possible paths between patches $i$ and $j$ (including single-step paths). | $PC(v) = \frac{\sum_{i=1}^n \sum_{j=1}^n a_i a_j p_{ij}}{A_L^2}$ | Landscape networks | 59 |

There is a wide range of functions that can accomplish this transformation. For example, if a_ij_ measures the flow of information between nodes *i* and *j*, the functions 1-a_ij_, exp(-a_ij_), 1/a_ij_, log(1/a_ij_) and log(a_ij_/(1-a_ij_)) are all found in the graph-theoretical literature^29–32^. Note that one must pay attention to the range of values that the edge weights can span, and to the values induced by the transformation – for example, one must avoid the use of negative edges (e.g., log transformation of values between 0 and 1), as these can greatly hamper the interpretation of the results. Finally, if the edge weights represent probabilities, one should account for the independence (or lack of independence) of the different edges.

## AVOIDING THE PITFALLS

When computing any shortest path-related measure, various decisions need to be made, and the wrong decision could lead to unexpected results, as we will illustrate in the following examples. We focus on the effects on Betweenness (BC) and Closeness centralities (CC) as these are the most commonly used centrality measures. In Figure 2 we summarize this decision process.

**Figure 2.**
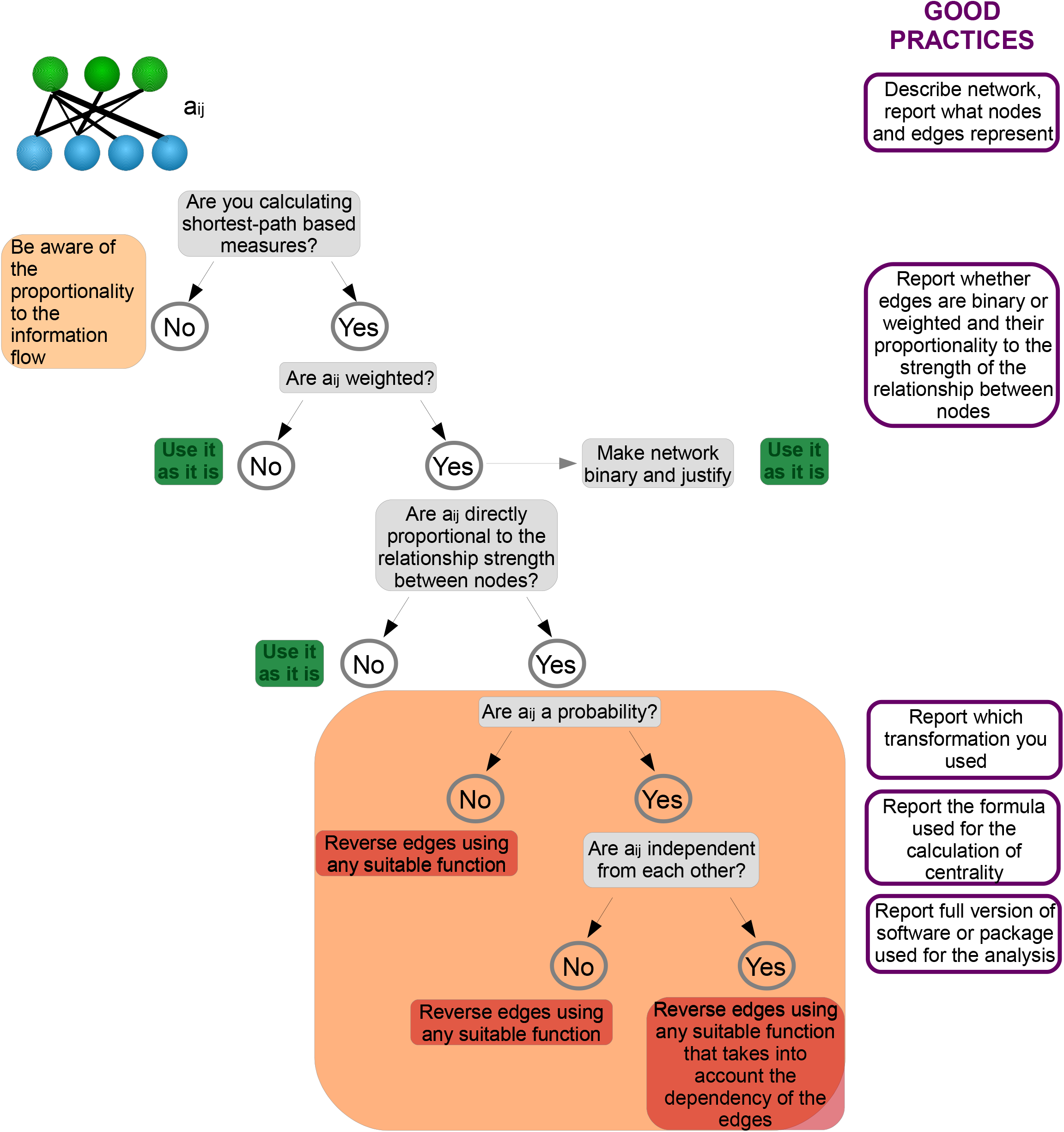
Scheme illustrating the step-by-step decision process for the calculation of shortest path-centrality measures in ecological networks and the 5-point guide of information require for good-practices.

### Binary or weighted?

The first methodological choice when computing shortest paths is whether to consider edge weights. Binary data can be highly informative: for example, it has been used to identify species fundamental niches^33^, and key species in pollination networks^18^. Nonetheless, studies must clearly state whether the analysis is performed on weighted or unweighted networks, and provide an ecological justification supporting either choice^34^. Seminal studies^35^ as well as recent ones^10^ analyzed unweighted versions of their data arguing that paths with fewer edges would inherently be stronger than those composed of multiple ones. In our analysis of the literature, we have found nine studies that, despite using weighted data for some calculations, revert to binary versions of the network for the calculation of centrality measures without providing a justification (Figure 1). Indeed, note that unipartite projections of bipartite networks result in weighted networks even when the original network is binary. Seven articles out of nine use binary unipartite projections without any justification. Although calculations in these are technically correct, one must be aware that the calculation of shortest paths and related measures in binary and weighted versions of the same network can lead to dramatically different conclusions.

To see how discarding the edge weights can significantly change the results of network analysis, let us start with a highly idealized example. In Figure 3a we show a network such that there are two possible paths between from Busan (South Korea) to Almería (Spain): one containing two edges (Busan-Chicago-Almería), and the other one three (Busan-Copenhagen-Marseille-Almería). If one considers only the binary data (implicitly assuming that the distance between any cities is the same), the shortest path from Busan to Almería would be crossing Chicago (one stopover vs. the two stopovers of the other possible path). However, if the distance between cities is considered, it can be easily seen that the path Busan-Copenhagen-Marseille-Almería is much shorter (~11000 vs. ~18000 Km).

**Figure 3.**
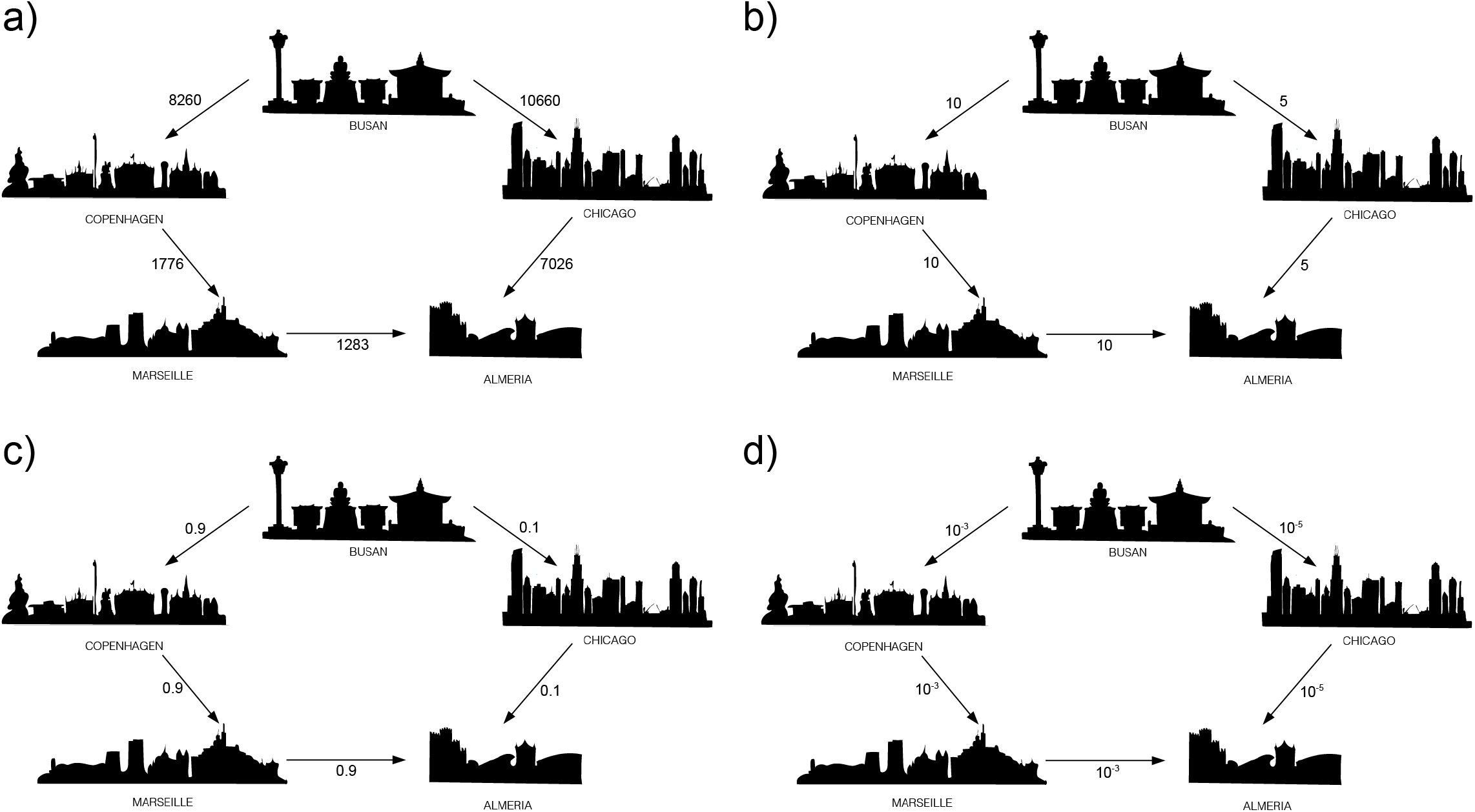
Toy networks depicting two possible paths to arrive from the city of Busan to the city of Almería. Edges connecting the different cities quantify distance in Km (a), or using different measurements of the movement of researchers between these cities (b-d; see text for explanation).

For an ecologically relevant example, we constructed a connectivity matrix for a hypothetical bird species living between 500 and 2000 m above sea-level, and with a typical habitat size of about 15 km^2^. For this purpose, the Global Relief Database ETOPO1^36^ data in the region we considered were coarse-grained to 15 km^2^ horizontal resolution, resulting in 787 habitat patches (Figure 4a). Dispersal probability between patches was calculated as p_ij_ = exp(-α d_ij_) following ref 35, where d_ij_ is the geographical distance between the patches boundaries, and α is a parameter chosen to be 0.03 in order to have a (hypothetical) median dispersal distance of 100 km. This probability can be stored in a connectivity matrix (see Supporting Information, Figure S1) and graph theory can be used to identify the habitat patches that have high Betweenness and Closeness centrality scores. Calculating BC and CC on binary and weighted versions of this dataset resulted in markedly different outcomes. None of the 20 habitat patches with highest BC and CC in the binary network (Figure 4b-c) match the ones obtained from the analysis of the weighted network (Figure 4d-e). Note that to calculate Betweenness and Closeness centralities in weighted data we first inverted the edge weights using log(1/p_ij_) (see next section for a detailed explanation).

**Figure 4.**
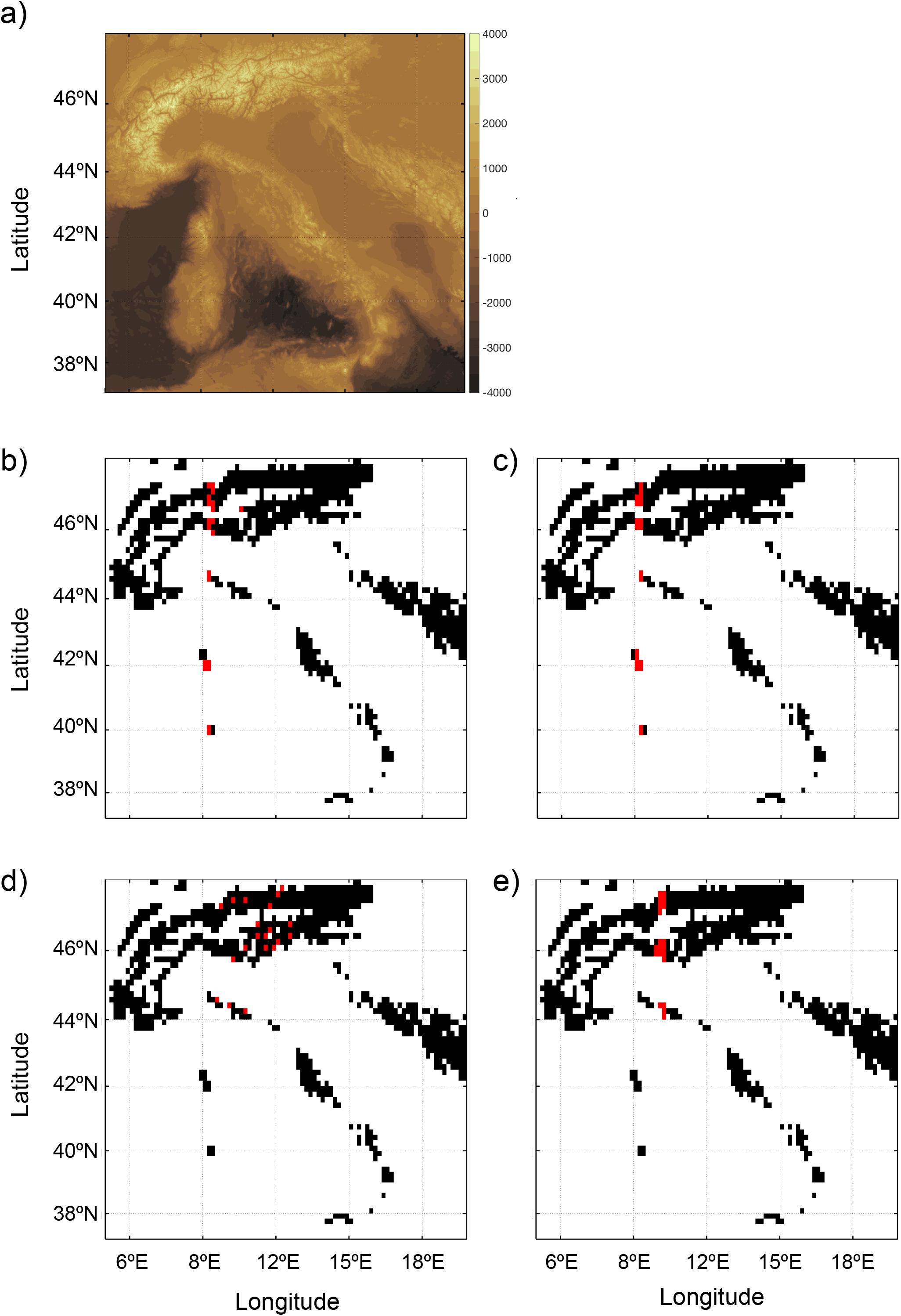
Land topography and ocean bathymetry (m) of the Italian region (data from the ETOPO1 Global Relief database; doi:10.7289/V5C8276M). A hypothetical bird species lives between 500 and 2000 m a.s.l., with a typical habitat size of about 15 km^2^ and a dispersal distance of 100km. a) Pixels in lighter tones denote patches of suitable habitat (colored in black in the following panels). After computing Betweenness (BC) and Closeness centrality (CC), we highlighted in red the 20 pixels with highest BC in the binary (b) and weighted network (c), and the 20 pixels with highest CC in the binary (d) and weighted network (e). Note the differences in the identification of the pixels between weighted and binary versions of the data.

### Modifying edge weights

The next question one should answer when computing shortest paths is whether edge weights are inversely proportional to the information flow between the nodes in the network (Figure 2). If this is the case, the shortest path between nodes can be calculated directly using the edge weights. If, on the other hand, edge weights are proportional to the flow of information, one must transform them before using shortest path algorithms. If one does not modify the edge weights, the shortest path algorithms will either fail (and identify the longest, rather than shortest path), or will be unable to identify a shortest path at all. We use the transformations 1/a_ij_ or log(1/a_ij_) for the edges (although for this operation there are several alternative options reviewed below).

For example, let us consider a small network of four primates sharing a certain number of parasites (Figure 5a). In order to detect the primate mediating the transmission of infectious diseases in this network, one could identify the primate displaying the largest number of parasites common to other primates – indicated by stronger edges. In this simple network it is easy to verify that most of the paths with the highest edge weight (largest number of shared parasites) pass through primate A. However, if BC and CC were calculated directly on unmodified edge weights, we would conclude that primate B is the key primate in this network (Figure 5a). If, on the other hand, we transform edge weights using the function 1/a_ij_, we correctly identify primate A as the primate with the highest BC and CC (Figure 5b).

**Figure 5.**
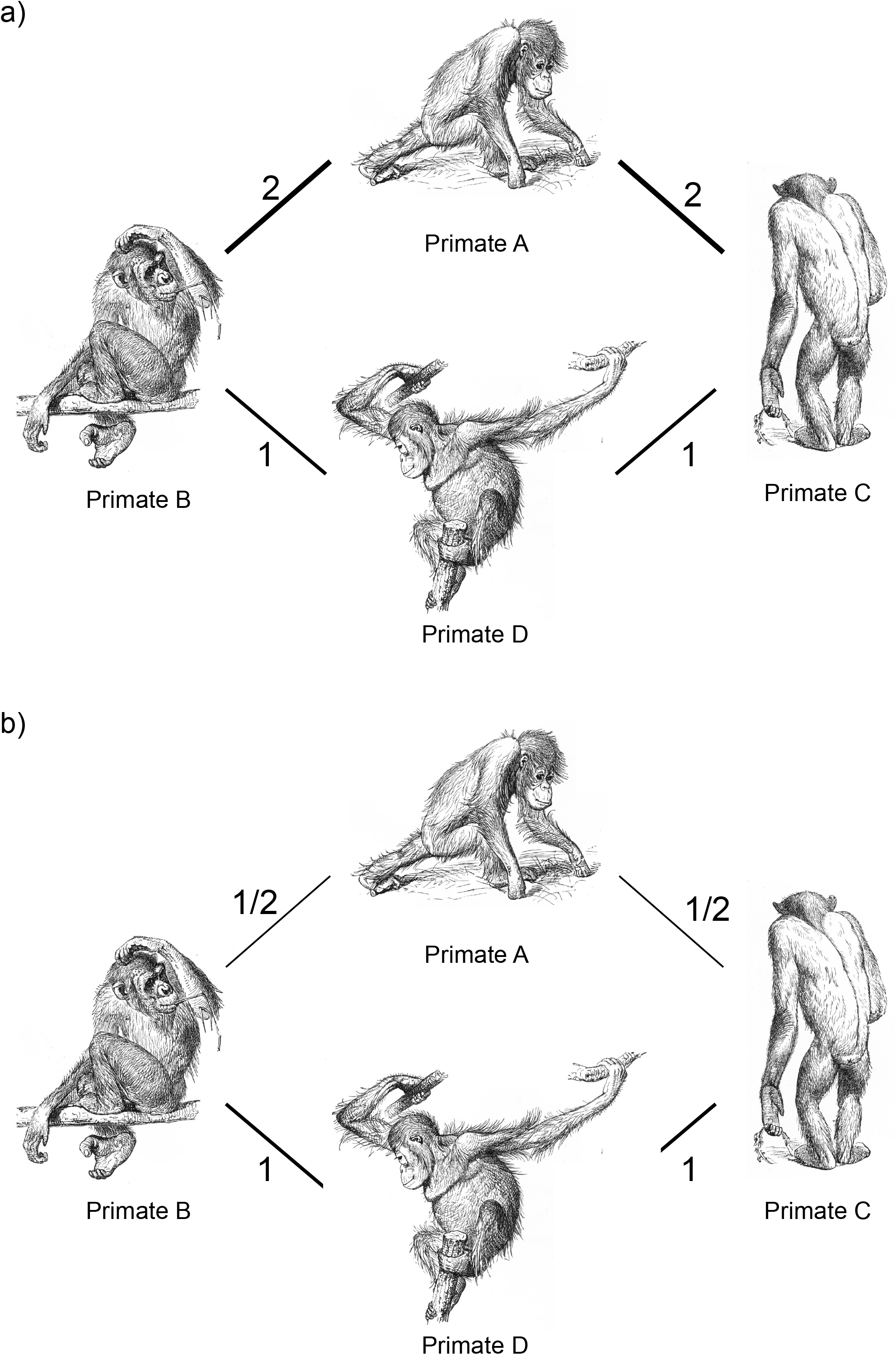
Toy network describing the social affinity between four primates. Edges quantify proportion of shared parasites. Drawings are public domain (https://commons.wikimedia.org/wiki/File:Primates-drawing.jpg).

Using a real-world case, the impact of not reversing edge weights can be illustrated using the Global Mammal Parasites Database (GMPD, https://parasites.nunn.lab.org) containing data on 542 primate species and their 750 parasites (ref 37 and references therein). To identify the species mediating the transmission of parasites (similarly to ref 38), a connectivity matrix linking primates that have been found to host the same parasite was built (see Supporting Information, Figure S2). Edge weights were defined as the number of shared parasites between any two species. Therefore, edge weight is directly proportional to the relationship strength between two species and, as in the previous example, the quantity should be transformed before calculating the shortest paths. In Figure 6a we show the lack of correlation between species rankings based on the centrality scores calculated on modified and unmodified edge weights. For this example, we use the inverse of the edge weights, e.g., 1/a_ij_. Interestingly, the BC scores calculated using the modified edge weights highlight only few species, one of which has by far the highest BC score. Instead, if we directly use the unmodified values, several more species have comparable BC scores. This is not surprising if we consider that, when using the raw weights, shortest paths pass through weak connections, which are likely to be numerous. Differences in ranks are substantial. For instance, among the top ten high-Betweenness species identified using modified weights, only one is also among the high-Betweenness species identified using raw weights (see Supporting Information, Table S1).

**Figure 6.**
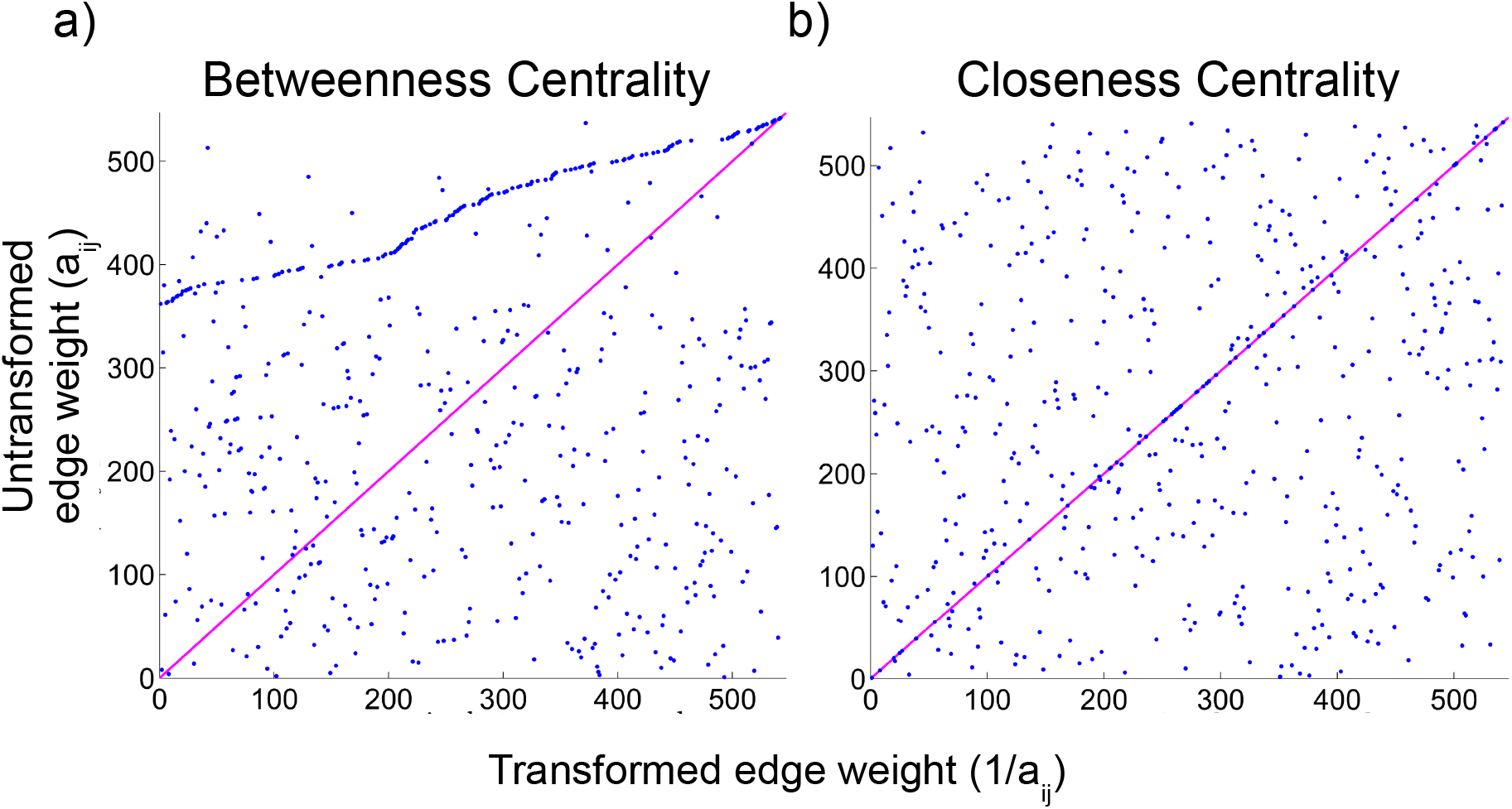
Scatter plot showing species ranks based on Betweenness (a) and Closeness centrality (b) values calculated on untransformed and inversed edge weights.

Likewise, the results from the Closeness-based rankings (Figure 6b) show that species rankings based on CC also differ significantly between modified and unmodified edge weights. The CC scores calculated on modified edge weights also support the importance of a handful of species (see Supporting Information, Table S1). Unlike with BC, nine of the top 10 high-Closeness species are the same ones when using the unmodified weights (also in Supporting Information) but the exact ranking differs between the two cases.

### Other modifying functions

#### Adding constants

When edges are directly proportional to the information flow, it is frequent practice to make them inversely proportional by subtracting their value from a theoretical maximum or some other meaningful constant^29,32^. For example, in the case of transfer probabilities (migration, mass, energy, networks), one could choose to subtract the edge weights a_ij_ from 1. However, the new edge weight 1-a_ij_ biases the calculation of the shortest paths towards the path with the lowest number of edges (because probabilities sum to one, nodes with many edges tend to have lower values).

Again, we will use the simple toy matrix presented in Figure 3, where we show a network connecting different cities. We then consider three different quantities to weight the edges that represent the movement of researchers between these cities. In the first case (Figure 3b), edge weights quantify the number of researchers who moved from one site to another. In this case, the path sustaining the largest “flow of researchers” between Busan and Almería is the three-steps path Busan-Copenhagen-Marseille-Almería (30 vs 10). However, applying a shortest path algorithm directly to this network would identify the two-step path as more important. One possible way to reverse the edge weight is to subtract the edge weights from a large (and in most instances arbitrary) constant C – de facto adding a constant to all edges. For example, if one chooses C = 100, now the largest edge weights (those representing largest flows) are the smallest and would hence be identified by the shortest-path centrality algorithm as more central. However, one can easily verify that such a transformation did not change the fact that the two-step path is the shortest between Busan and Almería (190 vs 270). The reason is that adding a constant to all the edge weights biases the shortest paths algorithm towards paths with fewer edges.

In Figure 3c we see that subtracting the edge weights from a theoretical maximum C (in the case of transfer probabilities of interaction frequencies, C = 1) makes the three-step path with higher information flow the shortest path (3x(1-0.9) vs 2x(1-0.1)). However, this transformation does not work in all cases. In fact, if the edge weights, as frequently occurs in ecological studies, span different orders of magnitude (e.g., Figure 3d), the three-step path will not be the shortest path anymore (3x(1-10^−3^) vs 2x(1-10^−5^)), as in Figure 3b. Furthermore, we note that this type of transformation, even when it is likely to work, cannot be used for all the edge weights. For example, it cannot be used for probabilities, as the values of log(1-a_ij_) are negative and, consequently, cannot be used to find shortest paths (see next section).

A real-world example of the effect of adding constants to the edge weights is provided in the Supporting Information (Figures S3, S4 and S5).

#### Negative weights and loops

Another way to reverse the edge weights is to reverse the sign of the weights (i.e., using −a_ij_). However, as shortest path algorithms seek to minimize the value of a path, they would keep looping closed paths (cycles) ad infinitum, without ever converging. It must be noted that there are alternative algorithms that can handle negative edges values (e.g., the Bellman-Ford-Moore algorithm^39^) cannot handle cycles. As cycles are essentially ubiquitous in ecological applications, edge weight transformations that result in negative values should therefore be avoided. As an example, consider a simple toy network depicting the carbon flow between different layers of a food chain (Figure 7a). In this case, given that the flow of carbon is directly proportional to the strength of the connection between two layers of the food chain, we need to transform the weights. However, if one uses −a_ij_ (Figure 7b), the shortest path algorithms would never converge, and would keep circling the loop. On the other hand, using another weight reversing function, such as 1/a_ij_, would correctly identify the 2^nd^ order consumers as key species pivoting the carbon flow in this network example (Figure 7c).

**Figure 7.**
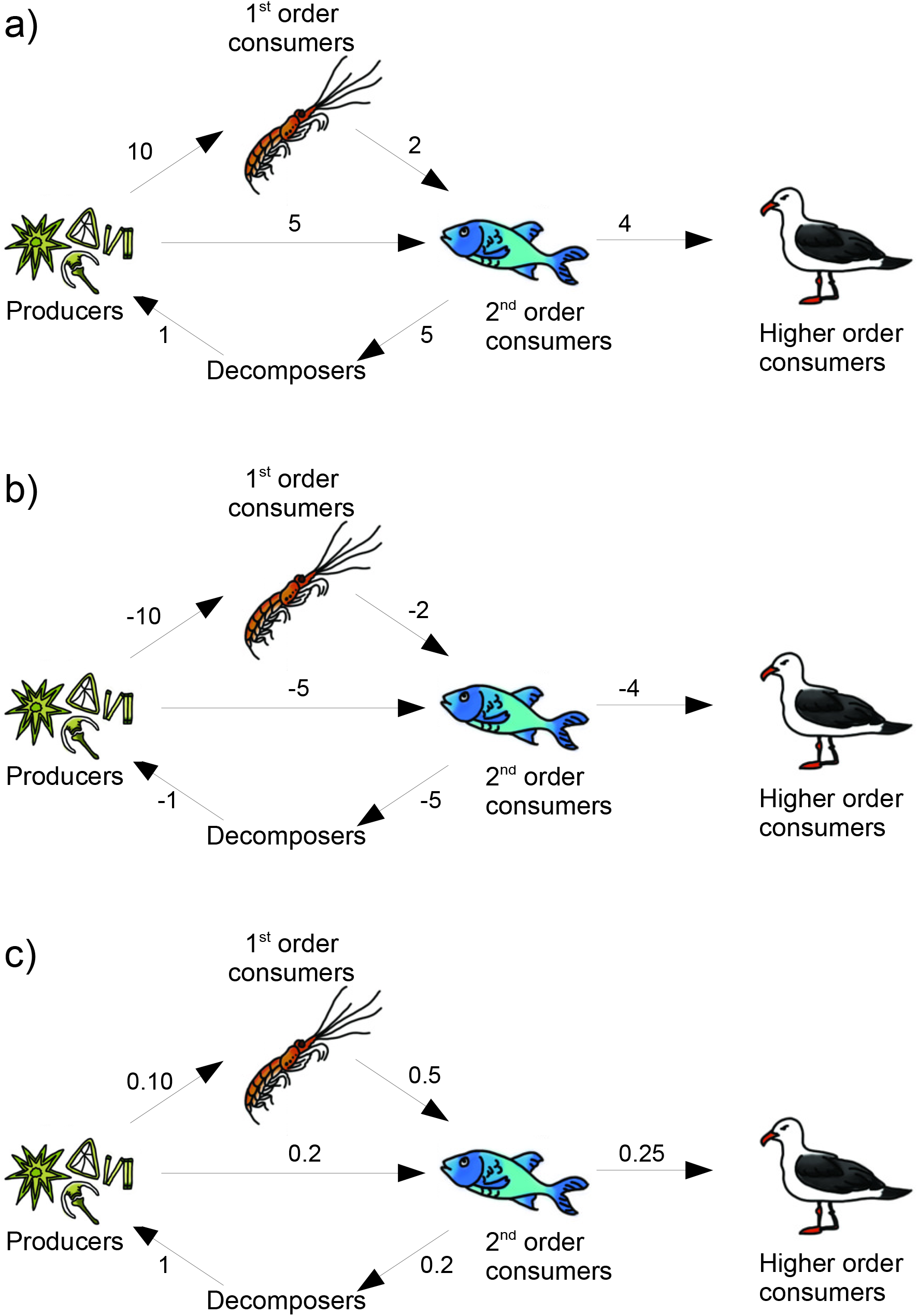
Toy network describing the carbon flow through a marine food web. Drawings by Siyavula Education under a CC BY 2.0 license (https://www.flickr.com/photos/121935927@N06/13578843423).

### Independence of probabilities

We should note an important aspect to consider when calculating the lengths of paths in networks: when edges represent probabilities, as for instance dispersal probabilities, we must question the independence of the edges in order to calculate meaningful values for the overall probability of the entire path. From a practical point of view, this means that when calculating the value of a path from node A to node C passing through node B (path ABC), we need to postulate that the path BC does not depend on the path used to reach B. When edges represent independent probabilities, the probability along a path containing multiple nodes is the product of the probabilities of all the paths linking the nodes. Interestingly, in the case of independent probability edges, converting edge weight a_ij_ into distance using log(1/a_ij_) nicely transform probability product along multiple nodes path into distances addition along this path, conserving weights relative contribution to the path and avoiding weights distortion in shortest paths algorithm. Without that edge transformation, the path ABC will be the sum of the probabilities of paths AB and BC, resulting in the identification of most improbable paths as those more central (see ref 24 for a detailed explanation, and Figure S6 for an example using the ETOPO1 dataset).

### One or all shortest paths?

In most networks, whether binary or weighted, there may be more than one shortest path connecting any two nodes. To account for this fact, an alternative definition of Betweenness centrality based on random walks^40^ and a generalization of node centrality that considers both edge weight and number when calculating centrality measures^32^ have been developed. Although the discussion of the pros and cons of the different formulations falls beyond the scope of this study, we encourage the reader to be aware that considering a single or all shortest paths may also introduce differences in the resulting centrality values, and the decision should hence also be reported. This is of particular importance when comparing centrality metrics with food chain length related metrics, where all paths may be considered.

## WIDENING THE PATH

Network analysis has been developing quite independently in different branches of ecology. However, dissemination between ecological disciplines and reproduction of published studies are being hampered, at least partially, by the lack of transparency when describing the methodologies used. Establishing a protocol for the analysis and reporting of calculations would ease these obstacles, and boost the use of centrality metrics for unconventional uses. For example, in a species-interaction network (where species are typically considered closer if they interact with higher frequencies), one could purposely choose to calculate shortest path-centrality measures without transforming the weights in order to study the effect of weak interactions across the network. For this reason, we urge all the researchers applying graph theory to ecological data to pay special attention when reporting their calculations, and, in particular, to provide a description of the network and edge weight they used.

Here, we provide a checklist of crucial methodological information that should always be reported (Figure 2). Following this guide ensures the study reports sufficient information to allow reproducibility, a quick understanding of the methods by readers from other fields, and that the decision process prior calculations is done sequentially.

1. *A clear definition of what nodes and edges represent.* Nodes and edges depict different entities and relationships in different ecological studies. Nodes may represent proteins, genes, individuals, populations, species, sites, etc., and edges may depict interactions of different kind, or movement measured in numerous ways. A clear definition of nodes and edges enables a faster and deeper understanding of the rationale and methodology of the analysis by readers from different disciplines.
2. *Are edges binary or weighted?* If edges are weighted, one needs to report the proportionality of the edge weight to the information flow between the nodes in order to evaluate whether edges need to be modified. In particular, one should ensure that there is no contradiction between the weights of a network and the interpretation of shortest paths.
3. *Report the eventual transformation applied to edge weight before the calculation of centrality measures*. Furthermore, carefully justify any conceptual reason to not transform edge weights, or to use the unweighted versions of weighted data. These decisions result in the identification of different central nodes or edges, and hence should be justified from an ecological perspective.
4. *Report the formula used for the calculation of the centrality measure, and whether it considers all shortest paths or only one*. To ensure reproducibility and a deeper understanding of what the results represent.
5. *Report the full version of the software or package used for the calculation of centrality*. To ensure reproducibility and to account for potential future updates in the algorithms used by different packages.

## CONCLUSIONS

Graph theory enables to achieve precious insights on ecological networks. For this reason, it has gained popularity in ecology and has developed quite independently in different disciplines, becoming a routine analysis in ecological studies. Our analysis of the literature evidenced that this familiarity is however associated to a lack of methodological rigor in the published studies. Indeed, by reading the methodological sections of a large portion of the published studies, we were not able to clearly ascertain what edges represented when centrality measures calculations were carried out. The increasing popularity of packages for the analysis of ecological networks will only boost the use of tools and methodologies researchers may be unfamiliar with. Using both theoretical and real-world case studies we showed that oversights in the methods and calculations can lead to radically different results. Hence it is fundamental to establish a code of good practices that guides researchers through the calculations, while ensuring the correct calculation of metrics across fields, aiding understanding from other fields and the reproducibility of results. For that reason, in this article we provide an overview of different methods to meaningfully calculate shortest paths and related centrality measures in ecological systems, and a checklist to ensure clear and sufficient reporting of such calculations. We hope that following the protocol we suggest will further increase the popularity of centrality measures in ecology, and, at the same time, guarantee the reproducibility of these studies.

## ACKNOWLEDGMENTS

This work was supported by IBS-R028-D1. AMMG is supported through a Marie Skłodowska-Curie Individual Fellowship (H2020-MSCA-IF-2015-704409), and thanks the Danish National Research Foundation for its support of the Center for Macroecology, Evolution and Climate (Grant number DNRF96). We thank Sonia Agüera-González (@immunosoni) for the illustrations in Figure 3 (https://www.behance.net/soniaguera6595).

## AUTHOR’S CONTRIBUTIONS

AC, AMMG, SA, AAP, KG & AMD designed the study; AC conducted the data analysis, AMMG performed the literature search and AC & AMMG performed the literature analysis and wrote the first draft of the manuscript. All authors contributed substantially to the drafts and gave final approval for publication.

## ADDITIONAL INFORMATION

### Competing interests

The authors declare no competing interests.

### Data availability

This article uses no data.

## Notes

#### Summary of Updates

Extended explanations in some sections and removed all referrals to the manuscript as a review

