## Supplementary Information 1 for "Ecological networks: Pursuing the shortest path, however narrow and crooked"

**Supporting Information 2.** Review of other potential pitfalls on shortest-path centrality calculations and additional figures.

**Figure S.1**. Connectivity matrix containing the dispersal probabilities between the habitat patches.

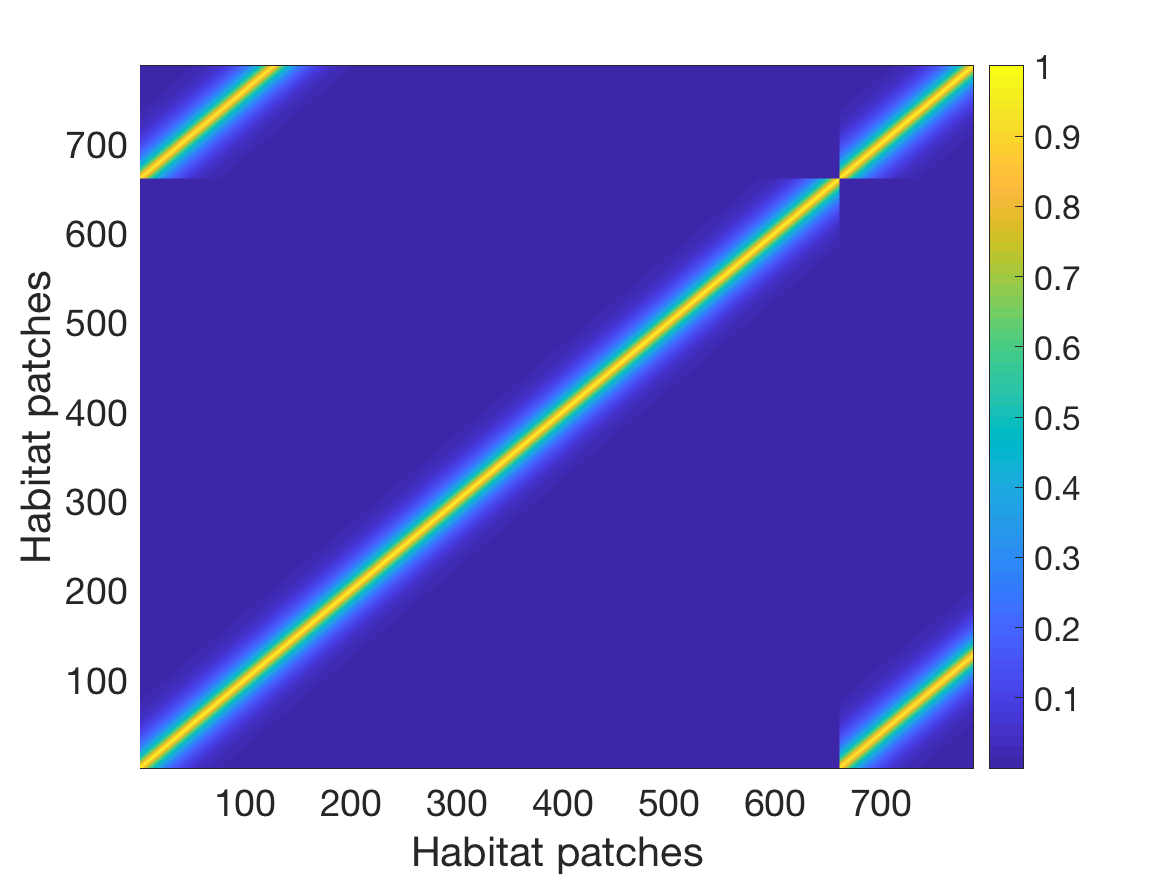

**Figure S.2.** Connectivity matrix and Top 10 species of the GMPD dataset containing the number of shared parasites.

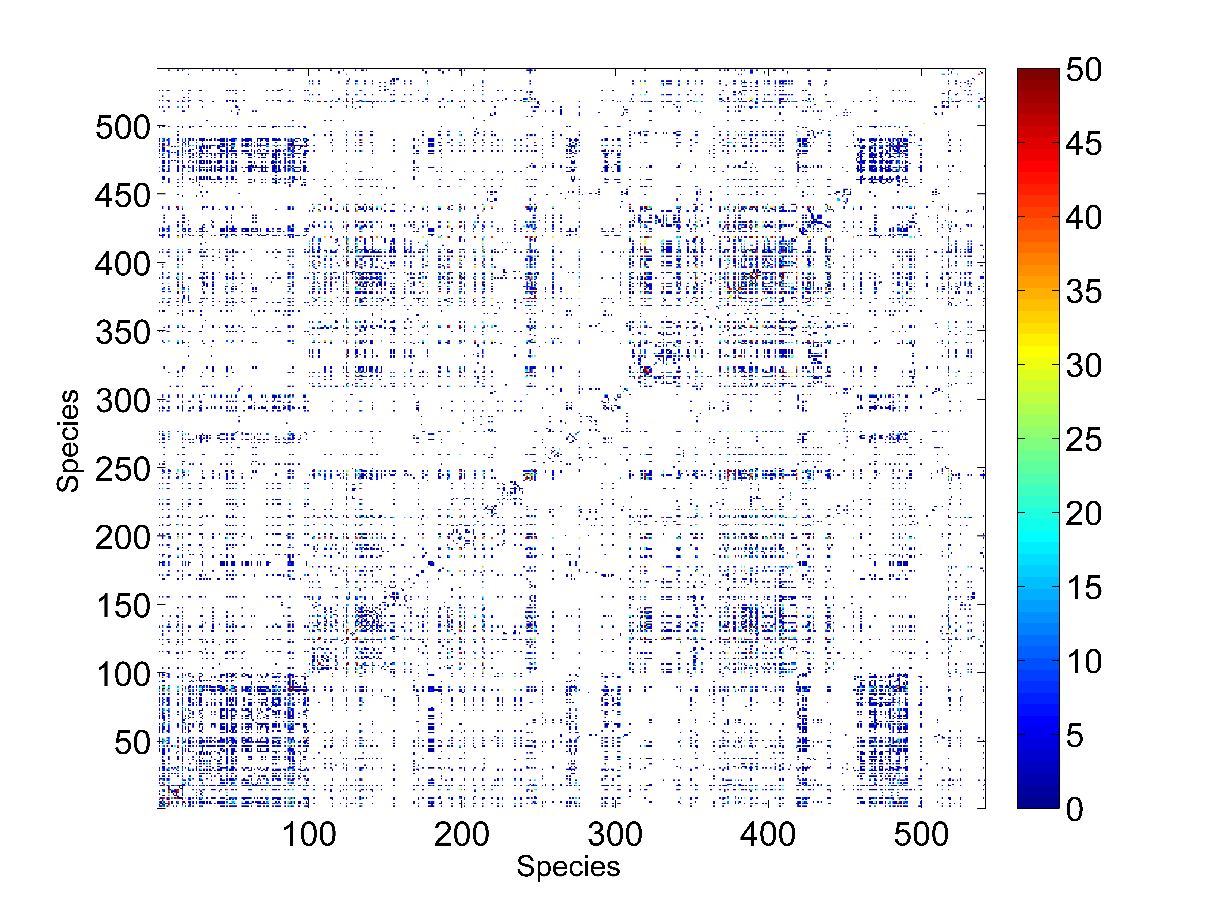

**Table S.1.** Top 10 species of the GMPD dataset identified on the base of betweenness and closeness values calculated with the a_ij_ and 1/a_ij_ edge weights.

| Betweenness ranking | | Closeness ranking | |
| --- | --- | --- | --- |
| 1/a_ij_ | a_ij_ | 1/a_ij_ | a_ij_ |
| *Pan troglodytes schweinfurthii* | *Papio sp.* | *Eulemur albifrons* | *Cercopithecus (Miopithecus) talapoin* |
| *Macaca fuscata* | *Cercopithecus nictitans* | *Eulemur macaco* | Crab-eating monkey |
| *Cercopithecus aethiops* | *Indri indri* | *Cercopithecus (Miopithecus) talapoin* | Taiwanese monkey |
| *Leontopithecus rosalia* | *Cercopithecus aethiops* | Crab-eating monkey | *Eulemur albifrons* |
| *Saimiri sciureus* | *Cercopithecus aethiops pygerythrus* | Taiwanese monkey | *Eulemur macaco* |
| *Cercopithecus ascanius schmidti* | *Pongo pygmaeus pygmaeus* | *Hylobates hoolock* | *Hylobates hoolock* |
| *Macaca fascicularis* | *Galago senegalensis* | *Hylobates lar entelloides* | *Hylobates lar entelloides* |
| *Papio ursinus* | *Macaca mulatta* | *Hylobates leucogenys* | *Hylobates leucogenys* |
| *Cebus apella* | *Saguinus fuscicollis* | *Macaca speciosa* | *Macaca speciosa* |
| *Lemur catta* | *Pithecia sp.* |  |  |

**Adding constants**

To show the biasing effect of adding constant on a real-world dataset, we exploited the data on a pollinator network (Robertson 1929) available from the Web of Life Ecological Database ([www.web-of-life.es](http://www.web-of-life.es/)). With the data we computed a connectivity matrix storing the interaction frequency between the different pollinators (which has a theoretical maximum of one if two pollinators were always found on all the plants):

**Figure S.3**. Connectivity matrix of the pollinators dataset storing the interaction frequency.

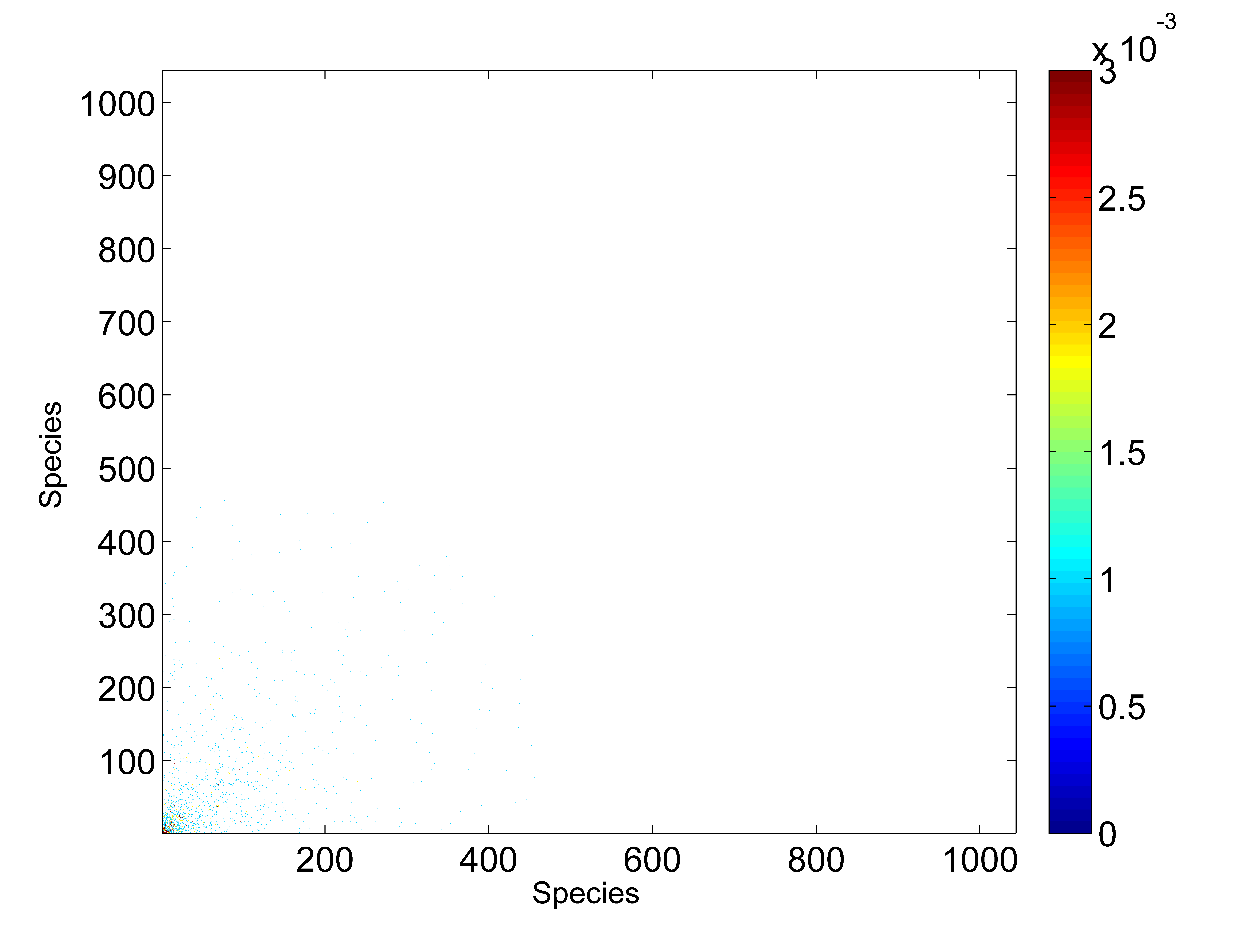

We can see that the matrix is very sparse. This will result in a limited portion of species to have a not null betweenness and closeness values.

Successively we computed the rakings based on betweenness and closeness using the two different edge weights a_ij_ and 1-a_ij_. The following figure shows the scatter plot between the rankings:

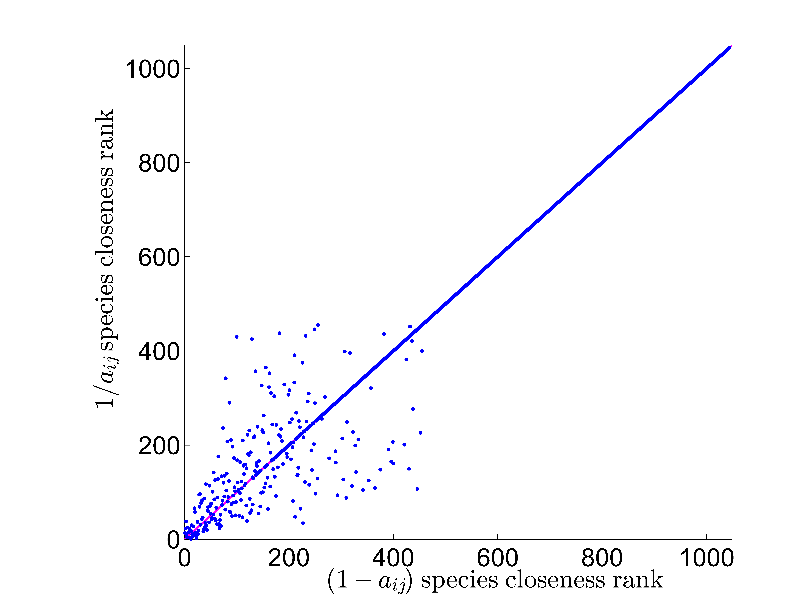
**Figure S.4.** Scatter plot of the betweenness (left) and the closeness-based (right) pollinators ranks using the 1/a_ij_ and 1-a_ij_ edge weights.

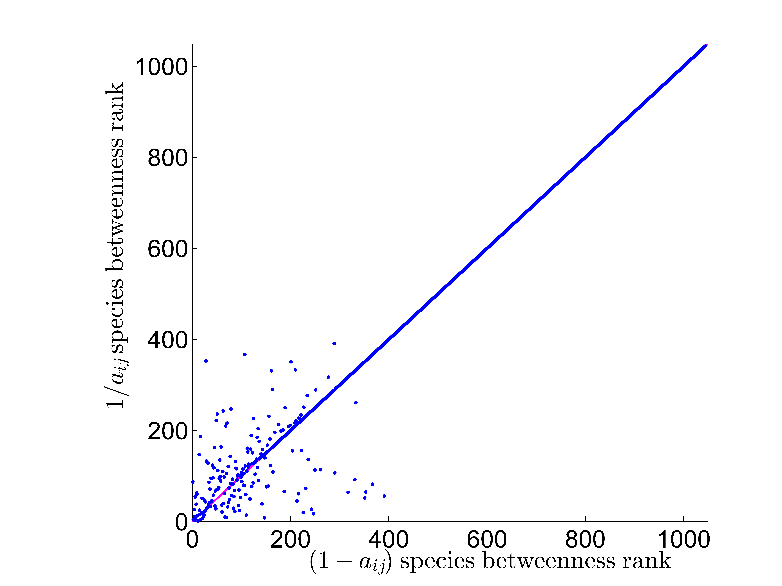
As it can be seen from the Figure, there is no clear correlation between the two ranks. Indeed, the dots fall on the bisecting line only for high ranks which correspond to zero betweenness and are determined by the alphabetical order with which the database was constructed. However, we can observe that there is some correlation between the ranks of high betweenness and closeness species. Indeed, if we look at the top 10 high-betweenness species, 7 are the same when using 1/a_ij_ or 1-a_ij_. However, the exact ranking differs:

| Betweenness ranking | |
| --- | --- |
| 1/a_ij_ | 1-a_ij_ |
| 1. 'Onthophagus pennsylvanicus' 2. 'Triepeolus donatus' 3. 'Anthocomus erichsoni' 4. 'Anoplius illinoensis' 5. 'Oscinis coxendix' 6. 'Spallanzania hesperidarum' 7. 'Hyalurgus johnsoni' 8. 'Euphorus mellipes' 9. 'Agapostemon texanus' 10. 'Andrena pruni' | 1. 'Onthophagus pennsylvanicus' 2. 'Hyalurgus johnsoni' 3. 'Andrena pruni' 4. 'Oscinis coxendix' 5. 'Anthrenus musaeorum' 6. 'Triepeolus donatus' 7. 'Scenopinus nubillipes' 8. 'Anthocomus erichsoni' 9. 'Agapostemon texanus' 10. 'Helophilus divisus' |

The same is true for the two closeness rankings (that share all the 10 species).

We also tested the effect of adding a constant to the edge weights on the ETOPO1 dataset. The following figure shows that top 20 betweenness and closeness patches identified using 1-a_ij_:

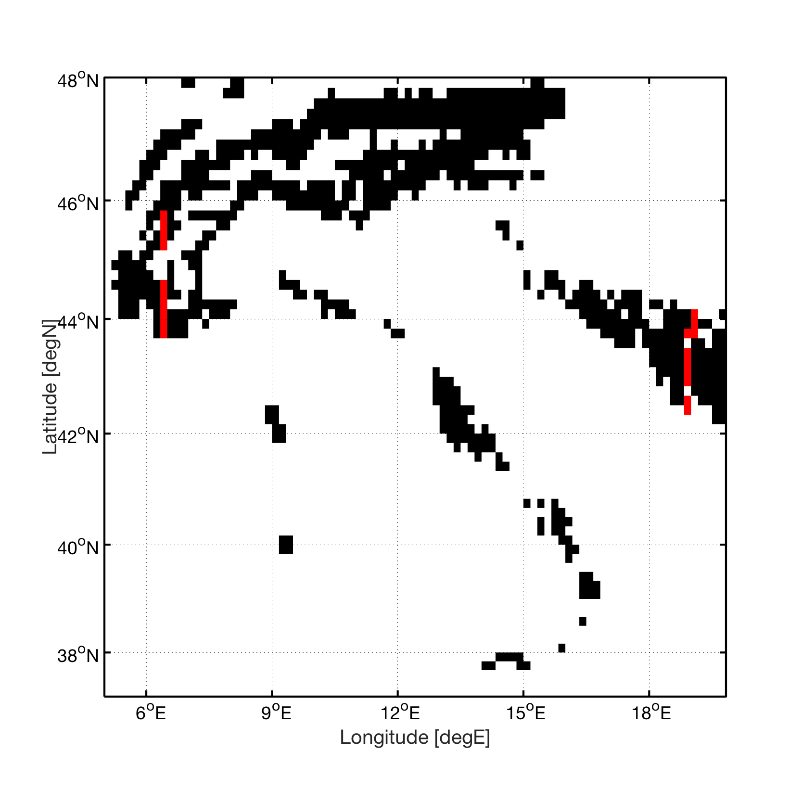

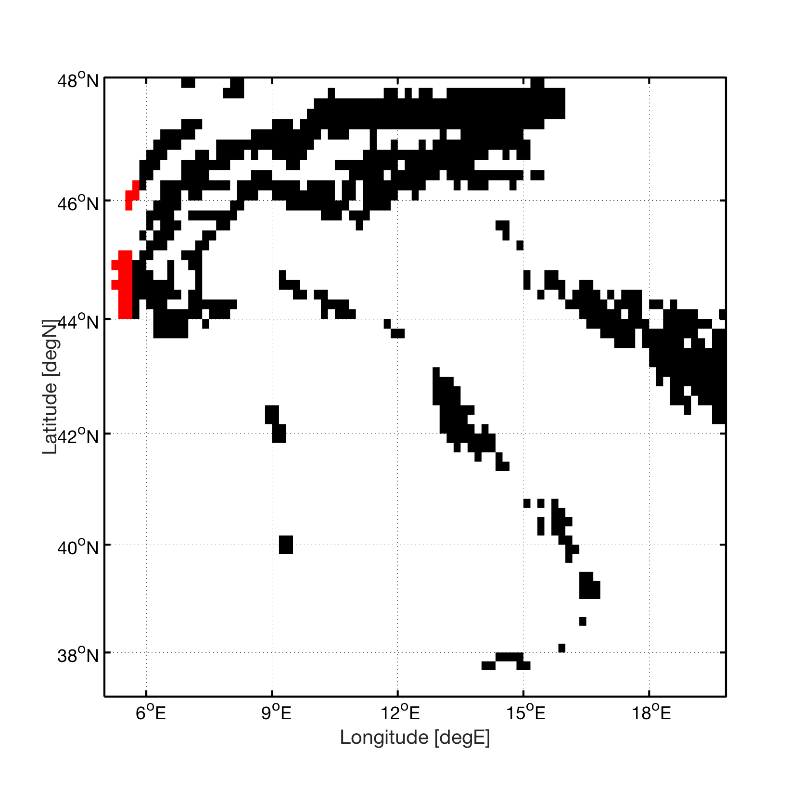

**Figure S.5.** Top 20 habitat patches based on betweenness ranking (left) and closeness ranking (right) using the 1-a_ij_ edge weights.

One can see that in this case the effects are much more drastic than in the pollinators dataset, as all the 20 habitat patches are different from the ones we identified when using log(1/a_ij_).

**Figure S.6.** Top 20 habitat patches based on betweenness ranking (left) and closeness ranking (right) using the 1/a_ij_ edge weights.

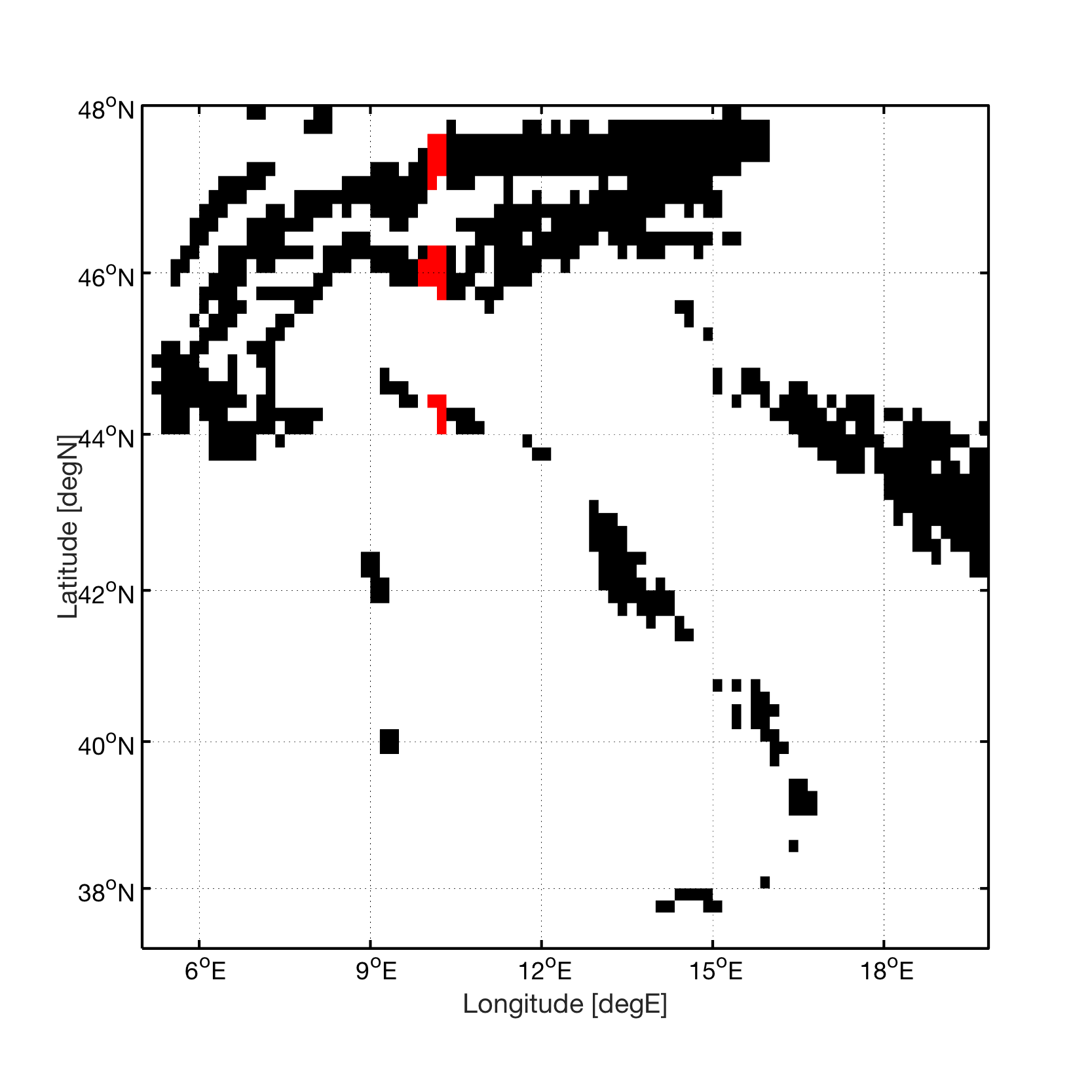

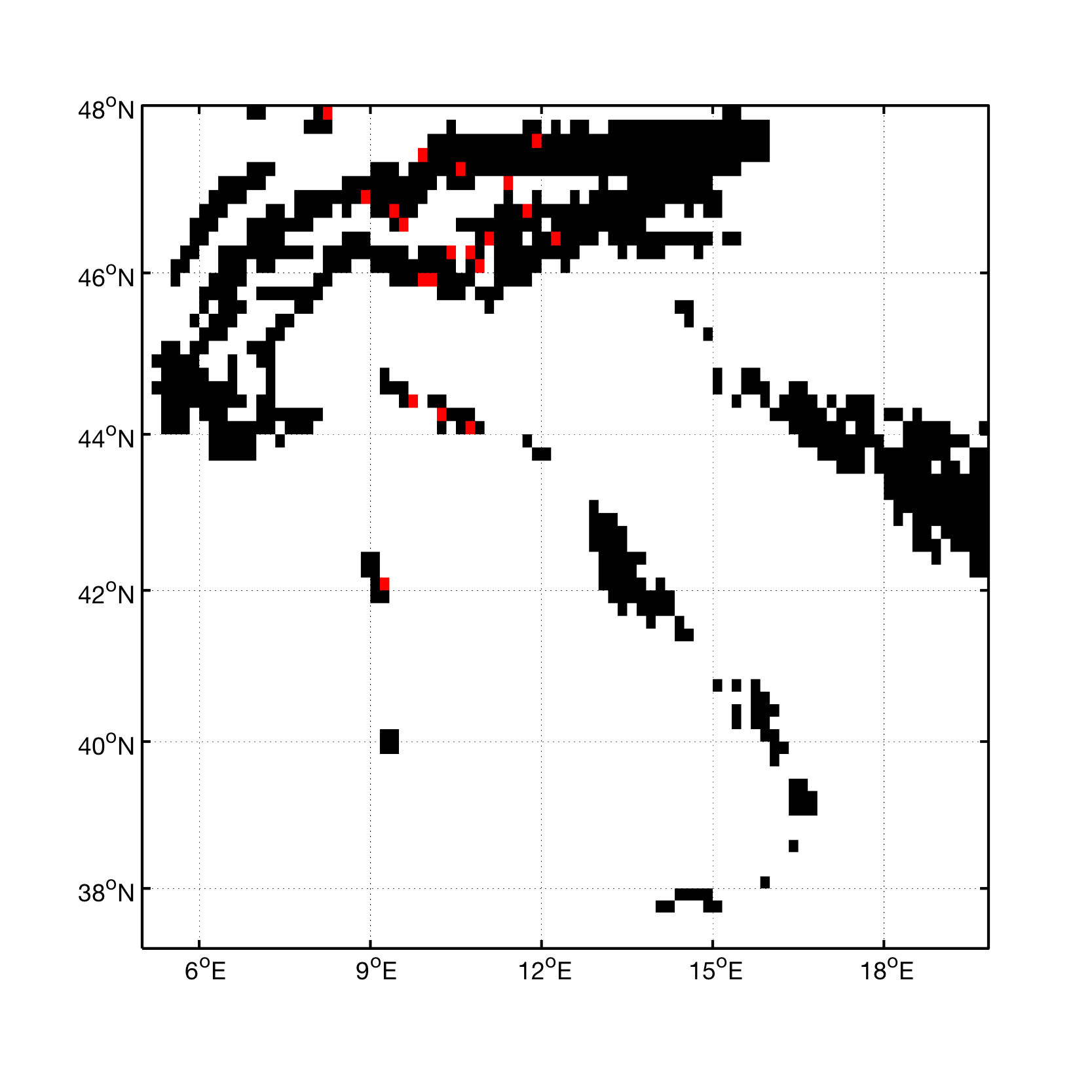
